# Central Amygdala Astrocyte Plasticity Underlies GABAergic Dysregulation in Ethanol Dependence

**DOI:** 10.1101/2024.06.11.598470

**Authors:** Todd B. Nentwig, J. Daniel Obray, Anna Kruyer, Erik T Wilkes, Dylan T. Vaughan, Michael D. Scofield, L. Judson Chandler

**Author notes:** **Corresponding Author:** L. Judson Chandler, PhD Department of Neuroscience Medical University of South Carolina 30 Courtenay St Charleston, SC 29425.

## Abstract

Dependence is a hallmark of alcohol use disorder characterized by excessive alcohol intake and withdrawal symptoms. The central nucleus of the amygdala (CeA) is a key brain structure underlying the synaptic and behavioral consequences of ethanol dependence. While accumulating evidence suggests that astrocytes regulate synaptic transmission and behavior, there is a limited understanding of the role astrocytes play in ethanol dependence. The present study used a combination of viral labeling, super resolution confocal microscopy, 3D image analysis, and slice electrophysiology to determine the effects of chronic intermittent ethanol (CIE) exposure on astrocyte plasticity in the CeA. During withdrawal from CIE exposure, we observed increased GABA transmission, an upregulation in astrocytic GAT3 levels, and an increased proximity of astrocyte processes near CeA synapses. Furthermore, GAT3 levels and synaptic proximity were positively associated with voluntary ethanol drinking in dependent rats. Slice electrophysiology confirmed that the upregulation in astrocytic GAT3 levels was functional, as CIE exposure unmasked a GAT3-sensitive tonic GABA current in the CeA. A causal role for astrocytic GAT3 in ethanol dependence was assessed using viral-mediated GAT3 overexpression and knockdown approaches. However, GAT3 knockdown or overexpression had no effect on somatic withdrawal symptoms, dependence-escalated ethanol intake, aversion-resistant drinking, or post-dependent ethanol drinking in male or female rats. Moreover, intra-CeA pharmacological inhibition of GAT3 also did not alter dependent ethanol drinking. Together, these findings indicate that ethanol dependence induces GABAergic dysregulation and astrocyte plasticity in the CeA. However, astrocytic GAT3 does not appear necessary for the drinking related phenotypes associated with dependence.

## Introduction

Alcohol misuse is a worldwide problem that represents a major health concern. In the United States alone, alcohol misuse is the 3^rd^ leading cause of preventable death and approximately 16 million Americans are diagnosed with an alcohol use disorder (AUD), the medical diagnosis for alcohol addiction[1]. AUD is defined as a chronic relapsing disorder that is characterized by compulsive alcohol seeking, loss of control over alcohol use, and the development of a negative emotional withdrawal state in the absence of alcohol[2, 3]. Alcohol dependence is defined by the emergence of an alcohol withdrawal syndrome when alcohol use is considerably reduced or suddenly stopped. Alcohol withdrawal consists of two categories of symptoms, somatic and psychological. Somatic symptoms may include central nervous system (CNS) hyperexcitability, elevated activation of the autonomic nervous system (eg, hyperarousal), and motor impairments (eg, rigidity, delirium tremens). Psychological symptoms primarily manifest as negative emotional states (eg, anxiety, depression, irritability, etc.) that can vary across individuals and promote maladaptive behavioral consequences such as risky decision-making, compulsive alcohol seeking and excessive alcohol intake[4]. The currently approved medications for AUD are marginally effective. Therefore, there is an urgent need to better understand the mechanisms contributing to AUD in order to develop improved therapeutic interventions [5].

The central nucleus of the amygdala (CeA) is a key brain structure underlying the neurobiology of ethanol dependence[6, 7]. Neuronal activity in the CeA modulates ethanol intake, withdrawal symptoms, and negative affective behaviors induced by chronic ethanol exposure[8, 9]. Moreover, chronic intermittent ethanol (CIE) exposure induces neuroadaptations in GABAergic transmission in the CeA that contribute to the behavioral consequences of dependence and withdrawal. For example, in vivo microdialysis studies have demonstrated that extracellular levels of GABA are increased in the CeA during withdrawal from chronic ethanol exposure[10, 11]. Furthermore, slice electrophysiology experiments have demonstrated that CIE-exposed rats exhibit increased spontaneous and evoked GABA_A_ receptor-mediated currents[6, 12] in the CeA. In addition, GABA_B_ receptor regulation of CeA neurotransmission is reduced in CIE-exposed rats, which likely contributes to elevated GABA tone and increased postsynaptic GABA_A_ receptor currents[13]. Together, these studies demonstrate that CeA activity contributes to ethanol intake and somatic withdrawal symptoms during dependence, and that chronic ethanol exposure dysregulates GABAergic transmission in the CeA. Despite the integral role of the CeA in ethanol dependence, most of our knowledge is based almost exclusively on a neuron-centric perspective, with little understanding of how non-neuronal cells may be involved. Astrocytes are a particular non-neuronal cell type of interest due to their ability to influence synaptic transmission and neuronal function.

In neuroscience, there was a longstanding view that astroglia were merely passive, support cells in the brain. However, within the past two decades, a wealth of research has demonstrated that astrocytes are active participants in neural processing and are essential for proper brain structure and function[14–16]. Astrocytes are abundant cells in the central nervous system with diverse functions that include regulation of blood flow, synapse development, neurotransmitter homeostasis, and synaptic function[14, 15, 17]. Accumulating evidence supports an essential role for astrocytes in regulating behavior due to their influence over synaptic transmission[18–21]. Astrocytes can directly regulate synaptic transmission via peripheral membrane processes that reside proximal to the presynaptic terminal, postsynaptic density, and extracellular matrix, which together comprise the tetrapartite synapse[22–24]. These perisynaptic astrocytic processes (PAPs) appear as fine, motile protrusions that ensheath synapses and express a variety of ion channels, G-protein coupled receptors, and neurotransmitter transporters[25–27]. PAPs influence synaptic transmission through two primary mechanisms, synaptic coverage and expression of neurotransmitter transporters[28, 29]. Due to their proximity to the synapse, PAPs are structurally positioned to bias the diffusion of neurotransmitters at the synapse. For example, PAPs preferentially contact the postsynaptic area and promote neurotransmitter diffusion towards presynaptic receptors on the axon terminal[30–33]. Alternatively, PAPs can encapsulate the presynaptic terminal and postsynaptic dendrite, restricting neurotransmitter spillover to neighboring synapses[31, 34]. Complementary to structural regulation of neurotransmitter diffusion, membrane-bound transporters provide PAPs with functional control of glutamate and GABA levels[35–38]. For example, astrocytes exclusively express the GABA transporter, GAT-3, which clears excess GABA from the perisynaptic space and shapes GABA receptor-mediated neurotransmission[38–46]. Together, the extent of synaptic coverage and surface expression of neurotransmitter transporters can permit or restrict neurotransmitter spillover, activation of extrasynaptic receptors, and overall synaptic efficacy[28].

The structural and functional mechanisms of synaptic regulation by PAPs are dysregulated in various pathophysiological conditions[39, 47–50]. Structural plasticity includes the reorganization of PAP synaptic proximity in response to synaptic activity or physiological states[16, 25, 27, 51–53]. Synaptic activity and consequent signaling cascades can facilitate protrusion or retraction of PAPs relative to the synapse, and thus alter synaptic events. Moreover, sustained alterations in synaptic proximity of astroglial processes can dictate long-term synaptic properties. For example, withdrawal from heroin self-administration is associated with reduced synaptic co-registration of the astroglial membrane and persistent neurotransmitter spillover[27, 51, 54–56]. Structural adaptations in PAP motility are often accompanied by functional plasticity that influences the overall impact on synaptic transmission. Functional PAP plasticity involves alterations in expression of membrane-bound neurotransmitter transporters that impact the efficacy and temporal properties of glutamate and GABA clearance[29, 37, 39, 47]. Transporter plasticity can manifest as changes in the overall transporter expression or surface vs. intracellular localization, which indicates the pool of transporters available for active use[57, 58]. For example, synaptic glutamate release or increases in astrocytic intracellular Ca^2+^ can alter the expression of surface-proximal glutamate transporter, GLT-1[58], or reduce expression of the GABA transporter, GAT-3, respectively[40]. Overall, structural and functional PAP plasticity is central to adaptive synaptic function and provides novel mechanisms amenable to therapeutic intervention. Given the dysregulation of CeA GABA signaling underlying ethanol dependence and the ability of astrocytes to shape GABAergic transmission, we hypothesized that astrocytes might be involved in key adaptations induced by chronic ethanol exposure. Therefore, the experiments described in the present study were designed to test the hypothesis that chronic ethanol exposure disrupts astrocyte-neurons interactions in the CeA, and that astrocytic adaptations will contribute to the negative behavioral consequences of ethanol dependence.

## Methods

### Animals

Adult male and female Sprague-Dawley rats (Envigo, Indianapolis, IN) were used for all experiments. All animal procedures were conducted with the approval of the Institutional Animal Care and Use Committee at the Medical University of South Carolina and adhered to the guidelines set forth by the National Research Council’s Guide for the Care and Use of Laboratory Animals. All methods and data are reported in accordance with ARRIVE guidelines (https://arriveguidelines.org) and recommendations.

### Surgical procedures

For astrocyte morphology experiments, astrocytes were labeled via injections of a membrane-localized GFP virus with an astrocyte promoter (AAV.GfaABC1D.LcK-GFP, Addgene #105598-AAV5) into the CeA (AP: -2.2-2.4, ML: 4.4, DV: 7.7-7.8 from skull surface). Viral vector approaches were used to manipulate astrocytic GAT3 protein expression. For overexpression of GAT3, animals received CeA injections of an AAV5 virus (packaged by University of South Carolina Viral Vector Core, Columbia, SC, USA) from the following plasmid: pZac2.1-GfaABC1D-GAT3-HA-p2A-mCherry (Addgene, #117722). GAT3 knockdown in astrocytes was achieved through delivery of an astrocyte-selective AAV5 serotype virus encoding a GAT3 shRNA[59] (pENN.AAV.HI.shR.GAT3.CMV.ZsGreen.SV40 plasmid obtained from Penn Vector Core; packaged by USC Viral Core). Viruses were injected at a rate of 2 nL/sec for a total of 1000 nL using a Hamilton syringe equipped to a micropump controller (World Precisions Instruments, Micro4, UMC4). Syringes were slowly withdrawn after a 5-minute diffusion period following the injection.

Bilateral guide cannulae (C315G; 26 gauge; 8mm length; P1 Technologies) were placed 2.0 mm above the CeA to enable pharmacological blockade of GAT3. Cannulae were surgically implanted 7 days prior to the start of CIE exposure and were kept patent by insertion of dummie cannulae. Rats received bilateral infusions (0.7 μL, 0.15 μL/min, 2-min diffusion) of the selective GAT3 blocker, SNAP-5114 (Tocris Cat # 1561), or vehicle 30 minutes prior to ethanol access. SNAP was dissolved in DMSO. All solutions were prepared fresh before use.

### Chronic intermittent ethanol vapor exposure and withdrawal drinking

Rats underwent the chronic ethanol exposure and withdrawal drinking paradigm previously described by our laboratory[60, 61]. This paradigm consists of several sequential phases: baseline ethanol drinking, chronic intermittent ethanol (CIE) exposure, ethanol drinking during withdrawal, and post-dependent ethanol drinking. In the baseline 2 bottle-choice (2BC) ethanol drinking phase, rats are allowed 24 hr access to voluntarily consume 20% ethanol and water in their home cage under an intermittent access schedule[62]. After 4 weeks of baseline drinking, rats are divided into control or ethanol vapor exposure groups to undergo CIE exposure. The CIE exposure procedure consists of 14 consecutive days of intermittent ethanol exposure by vapor inhalation. Rats are housed in ethanol vapor chambers for 14 hrs followed by 10 hrs out of the chambers, during which time they experience ethanol withdrawal. Control (AIR-exposed) rats underwent similar transport and handling procedures but were exposed to room air only. Tail-vein blood was collected periodically upon removal from the vapor chambers to determine blood ethanol concentrations (BECs) using an Analox alcohol analyzer (Analox Instruments, Atlanta, GA). Immediately prior to placing the rats back in the vapor chambers after the 10 hrs of withdrawal, somatic withdrawal signs were assessed using a behavioral rating scale[9, 63]. Withdrawal symptoms assessed included abnormal gait and tail stiffness, ventromedial limb retraction, and irritability to touch (includes vocalization). Each symptom was given a score of 0-2 (0 = no symptom, 1 = moderate, 2 = severe) and summated to provide a composite score (0-6) of withdrawal severity. For the present studies, intoxication levels were adjusted to achieve target withdrawal scores of 3-4. Following the 2 weeks of CIE exposure, animals transition to the CIE 2BC withdrawal drinking phase. During this phase, 3 days of CIE exposure were replaced with 3 days of intermittent 2BC withdrawal drinking. Withdrawal drinking consisted of 14 hr access to 20% ethanol and water in the homecage beginning 10 hrs into withdrawal following removal from vapor chambers. Each withdrawal drinking day was preceded and followed by a day of CIE exposure. Following 2 weeks of standard CIE 2BC drinking, rats were tested for aversion-resistant ethanol drinking using a standard quinine adulteration procedure[60, 64]. Increasing concentrations of quinine were added to the ethanol solution during individual CIE 2BC drinking sessions. Non-adulterated ethanol drinking sessions were interleaved between quinine-adulterated ethanol sessions. Ethanol intake during the non-adulterated sessions were averaged and served as the zero-concentration value. Following aversion-resistant drinking, the post-dependent 2BC ethanol drinking phase began, in which CIE exposure ceased and rats were instead provided daily 2BC access on the same 14 hr on/ 10 hr off schedule as CIE + 2BC drinking.

### immunohistochemistry

Animals were sacrificed via transcardial perfusion with 1X Phosphate Buffer (PB) and 4% paraformaldehyde (Electron Microscopy Sciences). Brains were extracted and kept in paraformaldehyde for up to 24 hrs before transfer into PBS. Brains were sectioned at 50 um using a vibrating blade microtome (Leica) and slices were stored in 1X PBS + 0.1% Sodium Azide. For immunofluorescence labeling, tissue sections were first permeabilized in 1X PBS + 2% Triton-X for 15 minutes. Tissue sections were then washed in blocking buffer (1X PBS + 0.2% Triton-X + 10% normal donkey serum) for 1 hour followed by overnight incubation at 4°C in blocking buffer plus primary antibodies including, mouse anti-synaptojanin1 (BD Biosciences #612248, 1:1000), rabbit anti-GAT3 (Abcam ab181783, 1:1000), guinea pig anti-GLT-1 (EMD Millipore, #ab1783), rabbit anti-GAT1 (Alomone Labs, #AGT-001, 1:1000), goat anti-GFP (Abcam, cat# ab5450 at 1:3000), or mouse-anti HA (Biolegend #901513, 1:1000). Following incubation with the primary antibody, the tissue was washed 3 times for 5 min each with 1X PBS + 0.2% Triton-X and then incubated overnight in blocking buffer containing Alexa-Fluor-conjugated secondary antibodies (Jackson ImmunoResearch Laboratories, 1:500). Upon completion of incubation with the secondary antibodies, tissue was washed 3 times in 1X PBS for 5 minutes each and stored in 1X PBS containing 0.1% Sodium Azide. Slices were mounted on standard, positively charged microscope slides and cover-slipped using ProLong Gold anti-fade mounting media (Invitrogen, cat# P36930).

### confocal microscopy and 3D image analysis

Astrocytes were imaged with a 63x oil immersion objective lens using a Zeiss LSM 880 confocal microscope with Airyscan[65]. Z-series images were collected using the Fast Airyscan module[66] at 1.8x digital zoom with a pinhole of 1 Airy unit, a 1024 x 1024 frame size, 0.2 um step size, and 4 frame averaging. Laser settings were identical across experimental conditions to enable proper comparison of protein expression. An average of 5-8 astrocytes were imaged per animal. Z-series were deconvolved using the airyscan processing function in the Zen Black software. Exported images were processed using Imaris image analysis software (Bitplane, Imaris) and a well-validated protocol for assessing colocalization of astrocytes with synaptic and membrane associated proteins[54, 67–69]. First, the Crop 3D function was performed to slightly crop images to the boundaries of the astrocyte. The surface function was then used to render a 3D model of the astrocyte surface based on the GFP signal intensity (**Figure 1A**). The 3D-rendered surface channel was masked over the original astrocyte channel prior to colocalization analysis. The Coloc module was then used to determine the co-registration between the astrocyte signal and either the astrocytic GABA transporter, GAT3 (**Figure 1B**), the synaptic marker, synaptojanin1 (SJ1; **Figure 1C**), or the triple co-registration between the astrocyte, GAT3, and SJ1 (**Figure 1D**). Co-registration of the GAT3 signal with the 3D rendered astrocyte volume is quantified to determine levels of astrocytic GAT3 expression. Co-registration of SJ1 signal with the astrocyte represents an index of synaptic adjacency of the astrocyte. Triple co-registration of astrocyte-SJ1-GAT3 reflects the proportion of GAT3-containing astrocyte processes that are proximal to the synapse. Percent co-registration is reported as normalized to the volume of the astrocyte surface.

**Figure 1.**
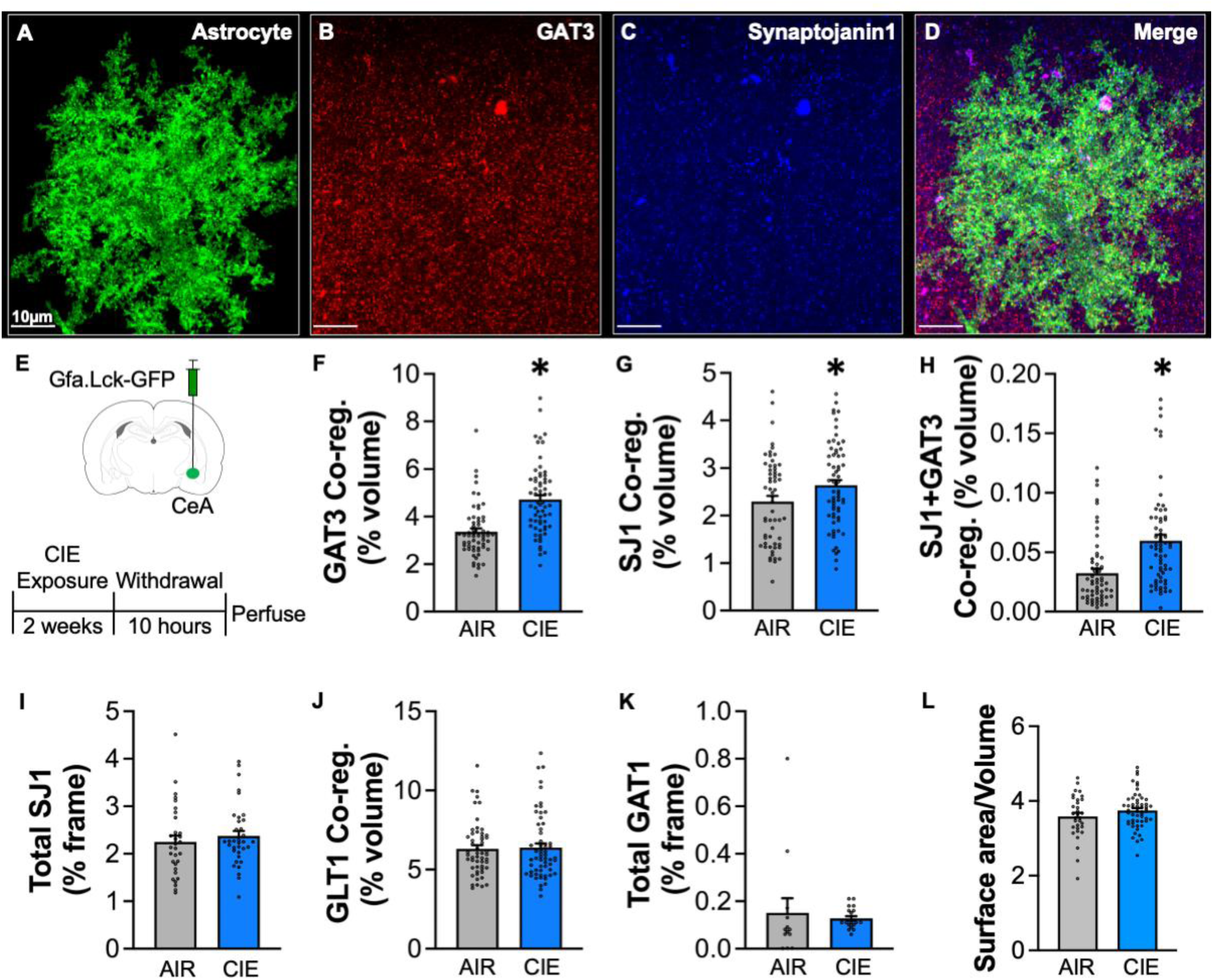
Central amygdala astrocytes upregulate GAT3 and increase synaptic proximity during withdrawal from ethanol dependence. **A**. A representative 3D-rendering of a CeA astrocyte labeled with an astrocyte-specific, membrane-localized, GFP reporter virus (AAV.Gfa.Lck-GFP). Representative immunolabeling for the astrocytic GABA transporter, GAT3, (**B**) the presynaptic marker, Synaptojanin1 (**C**), and their co-expression overlaid with the 3D-rendered astrocyte surface (**D**). **E**. Schematic depicting experimental approach. **F**. Average GAT3 co-registration. CIE-exposed rats displayed increased GAT3 co-registration, indicative of an upregulation in astrocytic GAT3 levels. **G**. Average Synaptojanin1 (SJ1) co-registration. CIE exposure induced an increase in SJ1 co-registration, suggesting an increase in astrocyte-synapse proximity. **H**. Average SJ1+GAT3 co-registration. CIE exposure induced an increase in SJ1+GAT3 co-registration, indicating an increase in the proportion of synapse-proximal, GAT3-expressing astrocyte processes. **I**. Average total SJ1 expression in CeA sections expressed as a percentage of the imaging frame volume. There was no difference in total SJ1 between AIR and CIE-exposed rats, suggesting CIE exposure does not alter overall SJ1 expression. **J**. Average Glutamate Transporter 1 (GLT1) co-registration. There was no difference in GLT1 co-registration between AIR and CIE-exposed rats. **K**. Average GABA Transporter 1 (GAT1) co-registration. There was no difference in GAT1 co-registration between AIR and CIE-exposed rats. **L**. Average astrocyte surface area normalized by volume. No differences in overall astrocyte size were observed between groups. *p < 0.05; n = 3-8 cells/4-8 rats per group.

### Slice electrophysiology

Brain slices containing the central amygdala (CeA) were obtained from AIR and CIE-exposed rats at 10 hours into withdrawal following the final ethanol vapor exposure. Rats were anesthetized using isoflurane and rapidly decapitated. The brain was promptly removed and placed into chilled sucrose based cutting solution consisting of (in mM): 200 sucrose, 1.9 KCl, 1.2 Na2HPO4, 33 NaHCO3, 10 D-glucose, 2.5 C5H9NO3S, 5 ascorbic acid, 10 MgCl2, and 0.5 CaCl2. Slices (300 µm thick) were prepared using a Leica VT1200 S vibratome and subsequently placed into aCSF consisting of (in mM): 92 NaCl, 2.5 KCl, 1.25 NaH2PO4, 30 NaHCO3, 25 D-glucose, 20 HEPES, 2.5 C5H9NO3S, 5 ascorbic acid, 10 MgCl2, and 0.5 CaCl2 maintained at 34 °C for 30 min before being incubated for an additional 30 min in room temperature aCSF. Following incubation, slices were transferred to a submerged recording chamber held at 34 °C and bathed in recording aCSF consisting of (in mM): 125 NaCl, 2.5 KCl, 25 NaHCO3, 10 D-glucose, 0.4 ascorbic acid, 1.3 MgCl2, and 2 CaCl2. The above solutions had an osmolarity of 300-310 mOsm, were pH adjusted (pH 7.3-7.43), and were continuously aerated with 5% CO2/95% O2.

Voltage clamp recordings were made using a Multiclamp 700B amplifier (Molecular Devices) connected to a Windows-PC running Axograph X software. Recording electrodes (2-4 MΩ resistance) were filled with a cesium chloride internal solution containing the following (in mM): 135 CsCl, 2 MgCl2, 10 HEPES, 1 EGTA, 4 NaCl, 2 Na-ATP, 0.3 Tris-GTP, 10 phosphocreatine, and 5 QX-314 Cl. The internal solution was pH adjusted (pH 7.3-7.43) with an osmolarity of approximately 285 mOsm. Neurons in the CeA were voltage clamped at -70 mV and all recordings were performed in the presence of CGP 55845 (1 µM), dl-APV (50 µM), and CNQX (10 µM). For measurement of inhibitory postsynaptic currents (IPSC), traces of spontaneous IPSCs (sIPSC) were recorded for 3 min after which tetrodotoxin (TTX, 1 µM) was added to the aCSF and mini-IPSCs (mIPSC) were recorded for an additional 3 min. For tonic currents, the baseline tonic current was measured for 2 min before determining the effect of GAT-3 inhibition on the tonic current by bath applying SNAP 5114 (20 µM). After the tonic current had stabilized following bath application of SNAP 5114 (minimum of 1.5 min), a combination of SNAP 5114 and Picrotoxin (100 µM) was perfused onto the slice and the tonic current was recorded for an additional 3 min. Recordings were then analyzed using a template matching algorithm to detect IPSCs and the mean holding potential to assess the effects of GAT-3 inhibition and GABAA receptor antagonism on the tonic current in neurons.

### Statistical analysis

Unless stated otherwise, all data was analyzed in GraphPad Prism 10 software using unpaired t-tests (with Welch’s correction when appropriate), linear regressions, and mixed effects ANOVAs, with post-hoc comparisons when applicable. Data are presented as mean ± sem, and effects were considered statistically significant at *p* ≤ 0.05.

## Results

### CEA astrocytes upregulate GAT3 and increase synaptic proximity during withdrawal from ethanol dependence

Rats received viral infusions into the CeA to label astrocytes before undergoing either AIR or CIE exposure and were sacrificed 10 hrs into withdrawal following the last exposure (**Figure 1E**). CIE-exposed rats exhibited increased GAT3 co-registration (**Figure 1F**; AIR: 3.35%, CIE: 4.17%; t_117.4_ = 5.864, p < 0.0001), suggesting that CIE exposure promotes upregulation of GAT3 expression on CeA. In addition, CIE-exposed rats had increased levels of SJ1 co-registration, (**Figure 1G**; AIR: 2.29%, CIE: 2.63%; t_122_ = 2.110, p = 0.03), further suggesting that CIE exposure induces an increase in proximity between synapses and astrocytes in the CeA. We then analyzed astrocyte-SJ1-GAT3 co-registration to determine whether the astrocyte processes that were proximal to synapses also contained GAT3. Relative to AIR controls, CIE-exposed rats exhibited an increase in co-registration of the astroglial membrane with SJ1 and GAT3 (**Figure 1H**; AIR: 0.032%, CIE: 0.059%; t_114.9_ = 4.352, p < 0.0001). These observations indicate that CIE exposure promoted protrusion of GAT3-containing astrocyte processes toward synapses in the CeA.

In tissue from a subset of the rats analyzed above, several controls were performed to assess specificity of the observed astrocytic adaptations. Total SJ1 puncta signal was quantified and normalized to the total volume of the imaging frame. No group differences were observed in total SJ1 (**Figure 1I**; AIR: 2.25%, CIE: 2.37%; t_66_ = 0.735, p = 0.46), indicating that CIE exposure didn’t alter overall expression of SJ1 in the CeA and that the increased co-registration (**Figure 1G**) was due to changes in astrocyte morphology.

Chronic ethanol exposure often results in widespread adaptations related to Glutamatergic and GABAergic neurotransmission[4, 11, 70]. Therefore, we assessed whether CIE exposure altered CeA expression of GLT-1, the astrocytic glutamate transporter 1[71], or the primarily neuronal GABA transporter, GAT1[41, 72]. This analysis revealed there were no differences between AIR and CIE-exposed rats for either GLT-1 co-registration (**Figure 1J**; AIR: 6.31 %, CIE: 6.39%; t_112_ = 0.2246, p = 0.8227) or total GAT1 expression (**Figure 1K**; AIR: 0.15%, CIE: 0.12%; t_12.59_ = 0.3633, p = 0.7224), suggesting that CIE exposure appears to preferentially alter expression of GAT3, but not GAT1 or GLT-1 transporters, in the CeA. Finally, to assess whether CIE exposure altered overall astrocyte size, the surface area of each astrocyte was normalized to its volume and quantified. Although, there was no group differences in astrocyte size between AIR and CIE-exposed rats (**Figure 1L**: AIR: 3.583, CIE: 3.745; t_84_ = 1.400, p = 0.1653).

### Chronic intermittent ethanol exposure-induced astrocyte plasticity persists beyond acute withdrawal

The next set of experiments sought to determine whether the CIE-induced changes in astrocyte plasticity persisted beyond the acute withdrawal phase following ethanol exposure (**Figure 2**). The approach for these studies were identical to the experiments in Figure 1 except that rats were sacrificed 7 days into withdrawal following their last day of CIE exposure (**Figure 2A**). This time point was chosen because it corresponded to the cessation of somatic withdrawal symptoms in CIE-exposed rats (**Figure 2B**). Similar to the 10 hr withdrawal time point, increased GAT3 co-registration was observed in CIE-exposed rats relative to AIR controls (**Figure 2C**; AIR: 3.04%, CIE: 3.77%; t_61.34_ = 3.034, p = 0.0035). In addition, CIE-exposed rats also exhibited a significant increase in SJ1 co-registration (**Figure 2D**; AIR: 2.50%, CIE: 3.68%; t_65_ = 3.199, p = 0.0021). However, there was no significant group differences in astrocyte-SJ1-GAT3 co-registration (data not shown). These results demonstrate that the CIE-induced increase in astrocytic GAT3 expression and astrocyte-synapse proximity extend well beyond the initial state of acute ethanol withdrawal.

**Figure 2.**
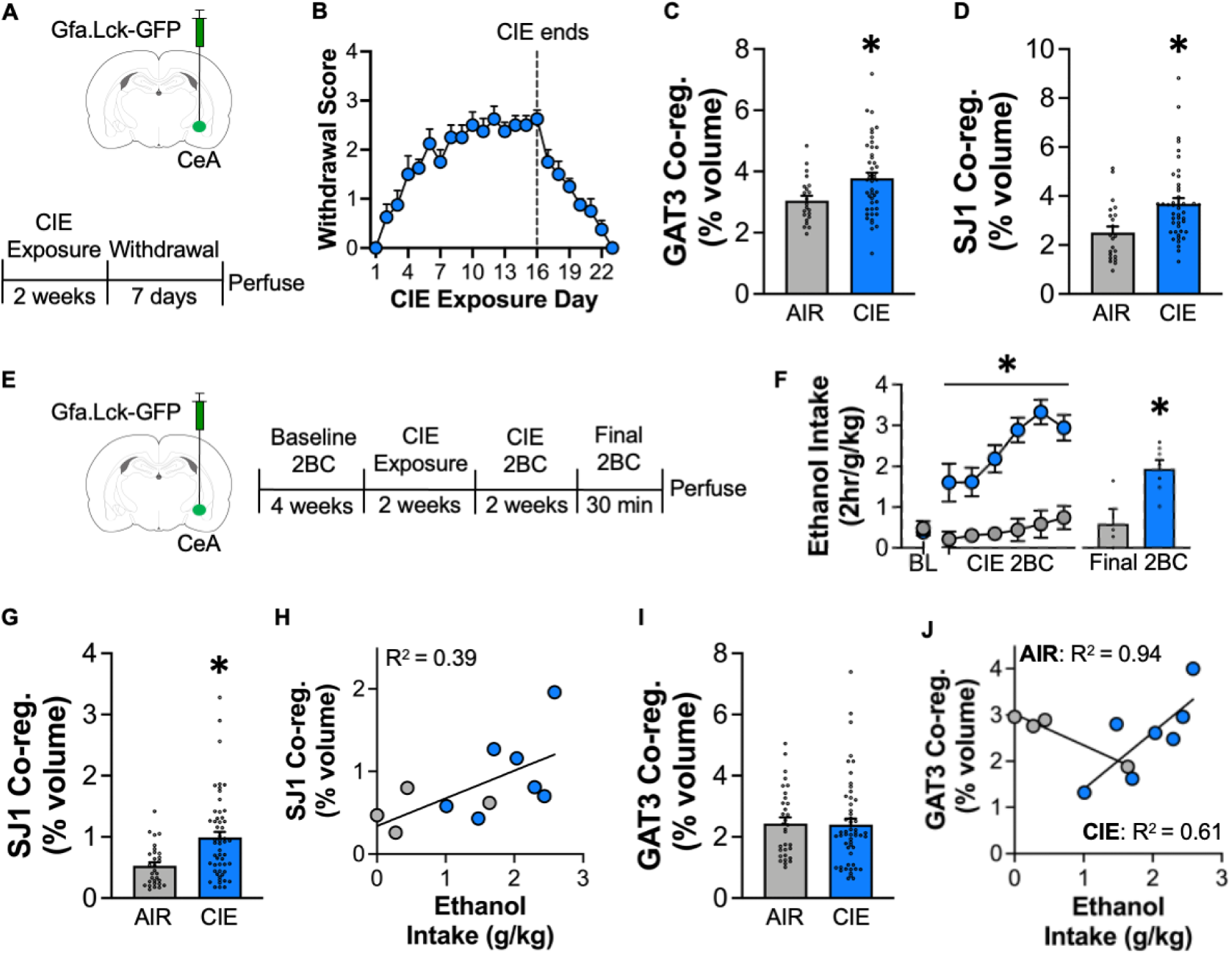
CeA Astrocyte Plasticity Persists Beyond Acute Withdrawal and Correlates with Voluntary Ethanol Intake. **A.** Schematic depicting experimental approach. **B.** Average withdrawal scores in CIE-exposed rats across the duration and cessation of CIE exposure. **C.** Average GAT3 co-registration. CIE-exposed rats displayed increased GAT3 co-registration, indicative of an upregulation in astrocytic GAT3 levels. **D.** Average Synaptojanin1 (SJ1) co-registration. CIE exposure induced an increase in SJ1 co-registration, suggesting an increase in astrocyte-synapse proximity. *p < 0.05; n = 6-8 cells/3-5 rats per group. **E.** Schematic depicting experimental timeline. **F.** Average 2 hr ethanol intake during baseline 2BC and CIE 2BC in AIR and CIE-exposed rats. CIE-exposed rats exhibited increased ethanol intake relative to their baseline (BL) intake (sessions 3-6 vs. BL) and AIR controls (all CIE 2BC sessions). Bar graph inset depicts 30 min ethanol intake during the Final 2BC session. CIE-exposed rats also consumed significantly more ethanol than AIR controls in this session. **G.** Average SJ1 co-registration in AIR and CIE-exposed rats following brief access to ethanol during withdrawal. CIE-exposed rats exhibited greater SJ1 co-registration compared to AIR controls. **H.** Linear regression analysis between SJ1 co-registration and ethanol intake in AIR and CIE-exposed rats. Overall, there was a positive relationship between ethanol intake and SJ1 co-registration. **I.** Average GAT3 co-registration. AIR and CIE-exposed rats displayed similar levels of GAT3 co-registration following voluntary ethanol intake. **J.** Linear regression analysis between GAT3 co-registration and ethanol intake. The relationship between GAT3 co-registration and ethanol intake was opposite in AIR and CIE-exposed rats. In AIR animals there was a negative relationship. Whereas CIE-exposed rats exhibited a positive association between ethanol intake and GAT3 co-registration, in which higher GAT3 co-registrations predicted greater levels of ethanol intake. *p < 0.05; n =7-9 cells/4-7 rats per group.

### Voluntary ethanol drinking during withdrawal modulates synaptic proximity and gat3 expression in CEA astrocytes

Dynamic changes in astrocyte morphology have been shown to occur rapidly in response to drug conditioned stimuli[29, 48, 54]. Therefore, the following experiment was designed to assess the effects of brief, voluntary ethanol drinking on measures of astrocyte plasticity. For these experiments, rats first underwent the CIE exposure and withdrawal drinking paradigm in order to establish stable escalation of ethanol intake during withdrawal (**Figure 2E**). The rats were then given 30-minute access to ethanol and water before sacrifice. CIE-exposed rats exhibited the typical escalation in ethanol intake relative to their baseline (BL) intake and AIR controls, as a mixed effects ANOVA revealed a significant treatment by drinking session interaction (F_(6, 60)_ = 7.756, *p* < 0.0001; **Figure 2F**). Post hoc comparisons indicated that CIE exposed rats had escalated ethanol intake in CIE 2BC sessions 3-6, relative to their BL intake, and all CIE 2BC session intake was greater than AIR controls (all *p*’s < 0.05). In addition, CIE-exposed rats also consumed significantly more ethanol than AIR controls in the 30-minute final 2BC session (AIR: 0.5881, CIE: 1.935; t_9_ = 3.445, p = 0.0073).

Similar to the 10 hour and 7-day withdrawal time points, a significant increase in SJ1-co-registration continued to be observed in the CIE-exposed rats compared to AIR controls (**Figure 2G**; AIR: 0.52%, CIE: 0.99%; t_80.94_ = 4.249, p = 0.0021). A simple linear regression analysis further revealed a significant positive relationship between ethanol intake and SJ1 co-registration (F_(1, 9)_ = 5.843, *p* = 0.0388, *R^2^* = 0.3937) (**Figure 2H**), suggesting that greater levels of astrocyte-synapse proximity were associated with greater levels of ethanol intake. In contrast, astrocytic GAT3 expression was not significantly different between AIR and CIE-exposed rats following brief ethanol access (**Figure 2I**; AIR: 2.43%, CIE: 2.40%; t_84_ = 0.1144, p = 0.9092). However, linear regression analysis revealed that GAT3 co-registration and ethanol intake were significantly associated (**Figure 2J**), but the direction of association was different in AIR and CIE-exposed rats, as indicated by significantly different slopes between groups (F_(1, 7)_ = 11.78, *p* = 0.0110). In AIR control rats, there was a significant negative relationship (F_(1, 2)_ = 37.22, *p* = 0.0258, *R^2^* = 0.9490) where higher ethanol intake was associated with lower GAT3 co-registration. In contrast, in CIE-exposed rats there was a significant positive relationship (F_(1, 5)_ = 7.948, *p* = 0.0371, *R^2^* = 0.6138), indicating that higher ethanol intake was associated with higher GAT3 co-registration. This suggests that rats with higher astrocytic GAT3 expression consume greater amounts of ethanol during withdrawal from CIE exposure. Examination of the triple co-registration between astrocyte-SJ1-GAT3 revealed no group differences or significant relationships with ethanol intake (data not shown). Overall, these results suggest that synaptic proximity and astrocytic GAT3 levels in the CeA are predictive of voluntary ethanol consumption during withdrawal from ethanol dependence.

### CIE exposure induces a GAT3-sensitive tonic current and increases GABAergic transmission in the CeA

The imaging data indicated an upregulation in astrocytic GAT3 expression in CIE-exposed rats. However, it is unclear whether this upregulation in GAT3 protein is functionally relevant to CeA physiology. Therefore, we tested whether CIE exposure produces changes in GAT3-sensitive tonic current and GABAergic tonic current in the CeA during withdrawal. Slice electrophysiology was performed on CeA slices obtained from rats during withdrawal following their final CIE exposure (**Figure 3A**). GAT3-sensitive tonic current and GABAergic tonic current were assessed by quantifying changes in holding current following bath application of the GAT3 blocker, SNAP-5114, and GABA-A receptor (GABAAR) antagonist, picrotoxin, respectively (**Figure 3B**). Compared to AIR controls, CIE-exposed rats exhibited a significant change in current during GAT3 blockade (**Figure 3C**; AIR: -4.048, CIE: -13.41; t_18_ = 2.235, p = 0.0383), suggesting that tonic GABA current is under greater regulation by GAT3 function in CIE-exposed rats. Furthermore, CIE-exposed rats also showed a significant change in current during picrotoxin application (**Figure 3D**; AIR: 5.989, CIE: 32.25; t_14.74_ = 3.269, p = 0.0053), suggesting that CIE-exposed rats exhibit a tonic GABA current during withdrawal. Together, these results suggest that CIE exposure promotes the emergence of a tonic GABA current in the CeA that is partially regulated by GAT3 function.

**Figure 3.**
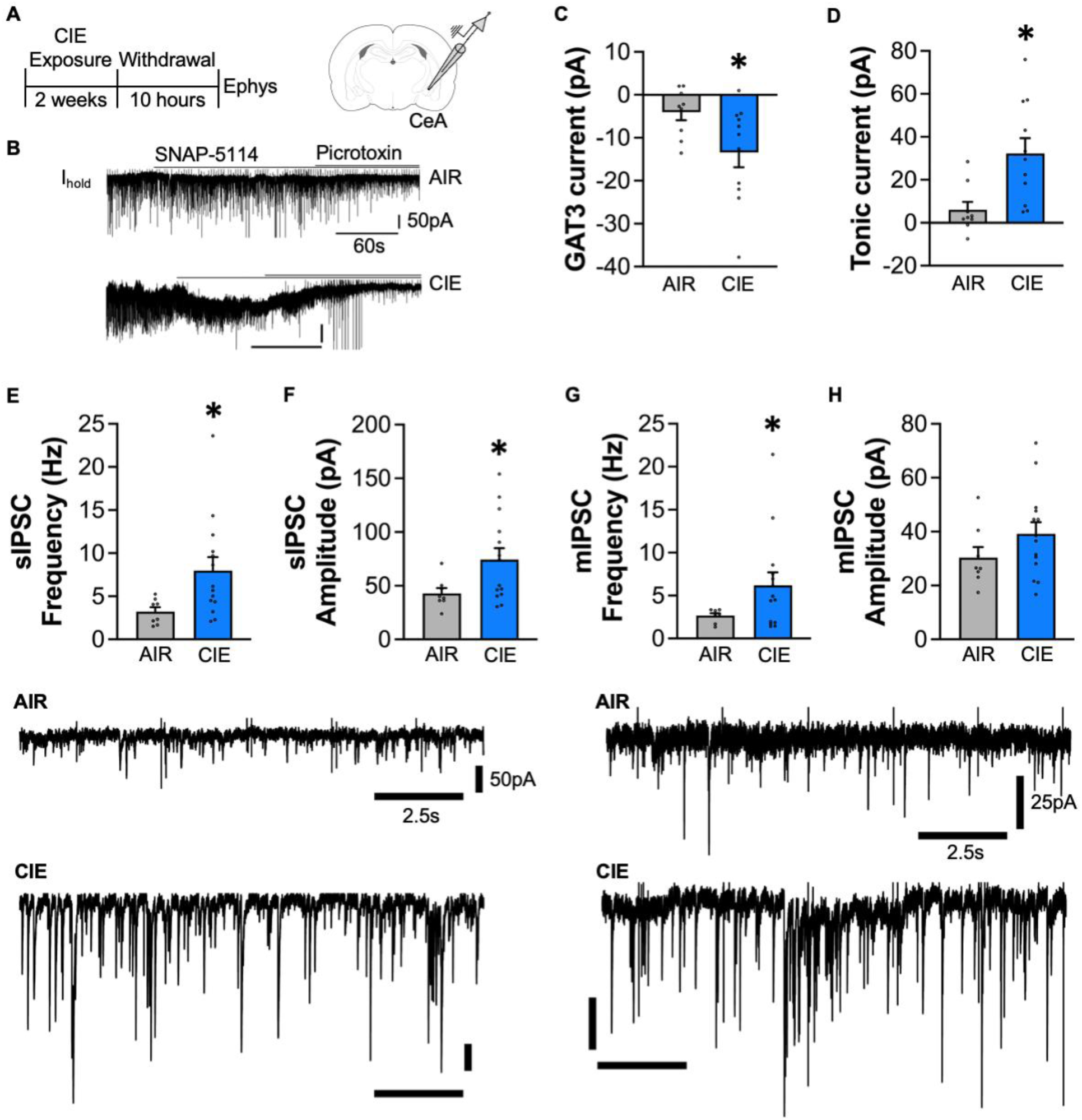
CIE exposure induces a GAT3-sensitive tonic current and increases GABAergic transmission in the CeA. **A.** Schematic depicting experimental approach. **B.** Representative traces from an AIR and CIE-exposed rat displaying the slice electrophysiology procedure for assessing GAT3-sensitive tonic current. In brief, a stable baseline holding current (I_hold_) is established prior to bath application of selective GAT3 blocker, SNAP-5114, followed by the GABA-A receptor antagonist Picrotoxin. **C.** Average GAT3 current in AIR and CIE-exposed rats during withdrawal from CIE exposure. CIE exposure induced a GAT3-sensitive tonic current not observed in AIR controls. **D.** Average tonic GABAergic current in AIR and CIE-exposed rats. CIE exposure induced a Picrotoxin-sensitive tonic current not observed in AIR controls. Average spontaneous inhibitory post-synaptic current (sIPSC) frequency **(E)** and amplitude **(F)**. Representative sIPSC traces from an AIR and CIE-exposed rat shown below. CIE exposure induced an increase in both sIPSC frequency and amplitude. Average miniature inhibitory post-synaptic current (mIPSC) frequency **(G)** and amplitude **(H)**. Representative mIPSC traces from an AIR and CIE-exposed rat shown below. CIE exposure induced an increase in mIPSC frequency, but not amplitude. *p < 0.05; n =1-5 cells/4-6 rats per group.

It is well documented that there is increased GABAergic transmission in the CeA during withdrawal from chronic ethanol exposure[6, 13, 73, 74]. Our working hypothesis is that the astrocyte plasticity observed in the present study is a homeostatic adaptation promoted by increased GABAergic transmission during withdrawal. Therefore, spontaneous inhibitory post synaptic currents (sIPSC) and miniature IPSC (mIPSC) were also collected to determine whether our model of CIE exposure produced increases in GABAergic transmission in the CeA. Compared, to AIR controls, CIE-exposed rats exhibited an increase in both sIPSC frequency (**Figure 3E**; AIR: 3.218, CIE: 7.993; t_15.67_ = 2.905, p = 0.0105) and amplitude (**Figure 3F**; AIR: 42.91, CIE: 74.28; t_17.54_ = 2.659, p = 0.0162). In addition, CIE exposed rats also had increased mIPSC frequency (**Figure 3G**; AIR: 2.676, CIE: 6.200; t_13.81_ = 2.307, p = 0.0371), but there was no group difference in mIPSC amplitude (**Figure 3H**; AIR: 30.32, CIE: 39.19; t_20_ = 1.358, p = 0.1895). The increase in sIPSC and mIPSC frequency indicates that there is increased GABA release in the CeA during withdrawal from ethanol dependence. Overall, the electrophysiology and imaging results indicate that following withdrawal from CIE exposure, there is increased GABAergic transmission and an increase in astrocytic GAT3 expression that functions to attenuate the GABA tonic current associated with extrasynaptic GABA spillover.

### Neither overexpression nor knockdown of GAT3 in CeA astrocytes alters dependence-escalated ethanol drinking

Thus far our results support the notion that CIE exposure induces increases in GABAergic transmission and spillover in the CeA that is associated with a homeostatic upregulation of astrocytic GAT3 expression and function. The next set of studies were designed to determine whether the CIE-induced adaptations in CeA astrocytes play a causal role in the behavioral consequences of ethanol dependence. The experimental approach involved assessing the effects of manipulating astrocytic GAT3 expression on dependence associated somatic withdrawal symptoms, escalated ethanol drinking, aversion-resistant drinking, and post-dependent drinking (**Figure 4**). Prior to initiation of the CIE exposure and withdrawal drinking paradigm, rats underwent surgery to transduce CeA astrocytes with a virus encoding a GAT3 shRNA or overexpression viral construct, or a control virus encoding GFP. Following recovery from surgery, rats began the baseline drinking phase of the CIE exposure and withdrawal drinking model (**Figure 4A**).

**Figure 4.**
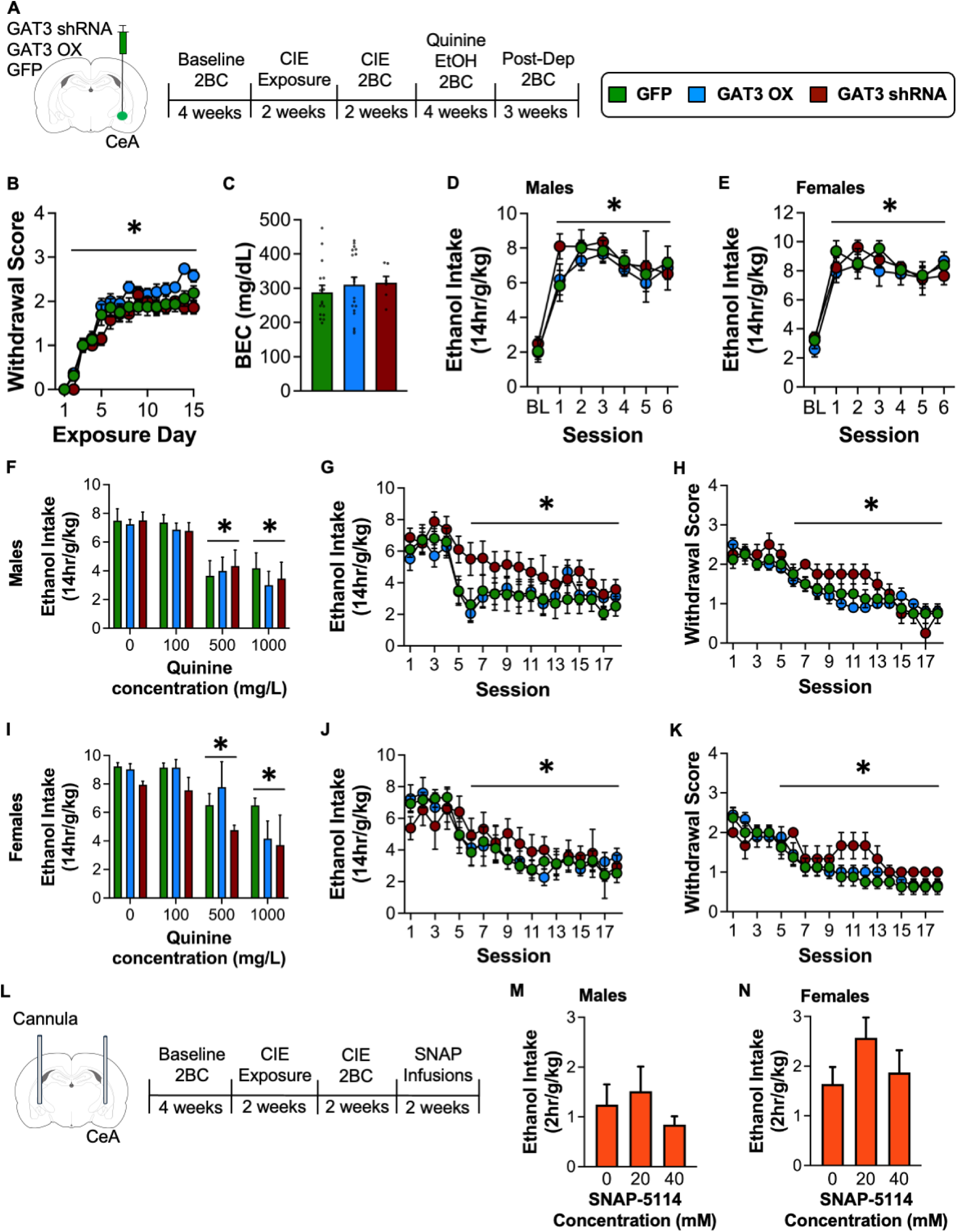
Effects of Virally Manipulating Astrocytic GAT3 Levels on Ethanol Dependence Phenotypes. **A.** Left: Schematic depicting experimental timeline. Right: Figure legend. **B.** Average withdrawal scores in CIE-exposed rats from control (GFP), GAT3 overexpression (GAT3 OX) and GAT3 knockdown (GAT3 shRNA) conditions. Ethanol vapor exposure induced somatic withdrawal symptoms similarly across all experimental conditions. Line above datapoints indicates that withdrawal symptoms were significantly elevated on exposure day 3 and onward, relative to day 1. **C.** Average blood ethanol concentrations (BEC) across CIE exposure in each viral condition. There was no significant difference in BECs between groups. Average 14 hr ethanol intake during CIE 2BC drinking in male **(D)** and female **(E)** rats. Asterisks indicate that, relative to Baseline 2BC drinking, all rats exhibited significantly escalated ethanol consumption throughout CIE 2BC drinking. Ethanol intake was not significantly different between virus conditions within either sex. Average 14 hr quinine-adulterated ethanol intake in male **(F)** and female **(I)** rats. The 500 and 1000 mg/L quinine concentration was sufficient to significantly suppress ethanol intake, relative to the 0 concentration, in male and female rats, regardless of virus condition. Average 14 hr post-dependent ethanol intake across daily drinking sessions in male **(G)** and female **(J)** rats. Regardless of virus condition, all rats showed a significant reduction in ethanol intake over time following cessation of ethanol vapor exposure. Average daily withdrawal scores during post-dependent 2BC drinking in male **(H)** and female **(K)** rats. All rats exhibited reduced withdrawal scores over time following cessation of ethanol vapor exposure and there were no significant differences between viral conditions in either sex. **L.** Schematic depicting timeline for the intra-CeA SNAP microinfusion experiment. Average 2hr ethanol intake in male **(M)** and female **(N)** rats following infusion of varying concentrations of the GAT3 inhibitor, SNAP-5114. There was no significant effect of intra-CeA SNAP administration on ethanol intake in either sex. *p < 0.05; n = 4-10/group.

Following baseline drinking all rats began CIE exposure to induce ethanol dependence. Withdrawal symptoms were assessed throughout the duration of CIE exposure in each virus condition (**Figure 4B**). A mixed effects ANOVA of withdrawal scores with virus condition and ethanol vapor exposure days as factors revealed a significant main effect of ethanol vapor exposure days (F_(7.643,295.4)_ = 72.97, *p <* 0.0001). Multiple comparisons following the main effect indicated that withdrawal scores on day 2 and beyond were significantly greater than the first day of exposure (all *p* values < 0.05), indicating somatic withdrawal symptoms increased over time and remained elevated similarly across each group. In addition, all groups achieved similar levels of intoxication throughout CIE exposure as there was no significant differences in blood ethanol concentrations (BEC) between groups (F_(2, 39)_ = 0.4210, *p* = 0.6594; **Figure 4C**). Following the CIE exposure phase, dependence-induced escalation of ethanol drinking was assessed in each group. A mixed effects ANOVA on 14 hr ethanol intake in male rats with virus condition and drinking session as factors revealed a significant main effect of session (F_(3.454, 61.03)_ = 37.92, *p <* 0.0001; **Figure 4D**). Multiple comparisons indicated that ethanol intake was significantly escalated, relative to baseline drinking, in all sessions, and similarly across all groups (all *p* values < 0.05). For the female drinking data (**Figure 4E**), a mixed effects analysis similarly revealed a main effect of session (F_(4.178, 71.03)_ = 26.94, *p <* 0.0001) with multiple comparisons indicating that ethanol intake was significantly escalated above baseline drinking in all sessions across each group (all *p* values < 0.05). Overall, these results indicate that knockdown or overexpression of GAT3 in CeA astrocytes does not appear to alter development of somatic withdrawal symptoms or dependence-escalated ethanol drinking in male or female rats.

### GAT3 overexpression and knockdown in CeA astrocytes does not influence quinine-resistant ethanol drinking

Prior research has implicated the CeA in the motivation to consume ethanol despite adverse consequences. For example, the CeA also mediates ethanol consumption when it is paired with an aversive footshock (punishment-resistant drinking) or adulterated with the bitter tastant quinine (quinine-resistant drinking)[59, 75, 76]. Thus, despite observing no differences in dependence-escalated drinking, we hypothesized that manipulating astrocytic GAT3 expression in the CeA may alter quinine-resistant ethanol drinking. Following the first two weeks of dependence-escalated drinking, rats continued 2BC drinking sessions, except their ethanol was now adulterated with various concentrations of quinine. A session with no quinine adulteration was always interleaved between a quinine session. A mixed effects ANOVA on 14 hr ethanol intake in males (**Figure 4F**) with virus condition and quinine concentration as factors revealed a main effect of concentration (F_(2.407, 43.32)_ = 24.62, *p <* 0.0001) but no effect of virus or interaction, suggesting that the quinine adulteration affected each group similarly. Multiple comparisons indicated that the 500 and 1000 mg/L concentration of quinine significantly reduced ethanol intake, relative to the 0 mg/L concentration (all *p* values < 0.05). In females, (**Figure 4I**), a mixed effects analysis similarly revealed a main effect of concentration (F_(1.625, 26.55)_ = 10.07, *p =* 0.0011) with multiple comparisons indicating that the 500 and 1000 mg/L quinine concentration significantly suppressed ethanol intake relative to the 0 mg/L concentration (all *p* values < 0.05). Therefore, neither knockdown or overexpression of GAT3 in CeA astrocytes altered the expression of quinine-resistant ethanol drinking during dependence in either male or female rats.

### GAT3 overexpression and knockdown in CeA astrocytes does not affect post-dependent ethanol drinking

Following testing for quinine-resistant drinking, rats proceeded to the post-dependent drinking phase. During this phase ethanol vapor exposure ends and rats transition to daily 2BC drinking. The post-dependent phase is thought to reflect a state in which acute somatic withdrawal symptoms dissipate but elevated ethanol intake and negative behavioral phenotypes still persist[77]. Ethanol intake and withdrawal scores were measured during this phase to assess how long elevated drinking and withdrawal symptoms persisted. A mixed effects ANOVA on 14 hr ethanol intake in male rats (**Figure 4G**), with virus condition and drinking session as factors, revealed a significant main effect of session (F_(3.756, 75.13)_ = 16.90, *p <* 0.0001) but no main effect of virus condition or interaction. Multiple comparisons indicated that ethanol consumption was significantly reduced at session six and onward, compared to the first session following the last ethanol vapor exposure (all *p* values < 0.05). The somatic withdrawal symptoms followed an identical pattern (**Figure 4H**) as a mixed effects ANOVA revealed a significant main effect of session (F_(5.028, 89.61)_ = 32.95, *p <* 0.0001) and multiple comparisons indicated that withdrawal symptoms were significantly reduced at session six and onward relative to session one (all *p* values < 0.05). This result indicates that following the cessation of vapor exposure, each group similarly reduced their level of ethanol consumption as their withdrawal symptoms dissipated.

Female rats exhibited a similar pattern to males across viral treatment conditions, as a mixed effects ANOVA of ethanol intake revealed a significant main effect of session (F_(5.103, 120.1)_ = 31.38, *p <* 0.0001; **Figure 4J**). Multiple comparisons indicated that ethanol intake was significantly reduced by session six and onward, compared to session one (all *p* values < 0.05). The withdrawal symptoms followed a similar temporal trajectory (**Figure 4J**) as a mixed effects ANOVA revealed a significant main effect of session (F_(5.174, 87.66)_ = 23.86, *p <* 0.0001) and multiple comparisons indicated that withdrawal symptoms were significantly reduced at session five and onward, relative to session one (all *p* values < 0.05). This result suggests that, like males, female rats reduce their ethanol consumption as their withdrawal symptoms abate.

### Pharmacological blockade of GAT3 in the CeA does not alter dependence-escalated drinking

Our viral-mediated GAT3 manipulations strongly suggest that GAT3 levels in CeA astrocytes are not necessary or sufficient for several dependence-related drinking phenotypes. However, it is possible that compensatory mechanisms during long-term viral mediated knockdown and overexpression may have obscured potential effects. Therefore, the next set of studies took a complementary pharmacological approach to inhibit GAT3 in the CeA of dependent rats via microinfusions of the selective GAT3 blocker, SNAP-5114 (**Figure 4L**). Following induction of dependence and stable escalation of ethanol drinking, rats received either vehicle or SNAP microinfusions into the CeA prior to CIE 2BC drinking sessions, and ethanol intake was then assessed 2 hours following bottle onset. In males, a mixed effects ANOVA revealed no significant effect of SNAP administration on ethanol intake (F_(1.123, 4.493)_ = 0.6058, *p* = 0.4944; **Figure 4M**). Similarly, SNAP had no effect in females (F_(1.984, 12.89)_ = 2.449, *p* = 0.1257; **Figure 4N)**. Taken together with the studies of viral manipulation of GAT3 expression, these data that despite being modulated by ethanol dependence, GAT3 in CeA astrocytes appears dispensable for a variety of dependence associated drinking phenotypes.

## Discussion

The experiments performed in the present study describe the effects of chronic alcohol exposure on astrocyte-neuron interactions in the CeA in rats. In addition, the effects of manipulating GAT3 in CeA astrocytes on the behavioral consequences of ethanol dependence were also examined. Following either acute or protracted withdrawal, dependent rats exhibit increased astrocyte-synapse proximity, increased expression of astrocytic GAT3, and increased synaptic proximity of GAT3-containing astrocytic processes (only during acute withdrawal). Additionally, voluntary ethanol intake during withdrawal was associated with the degree of synaptic proximity and GAT3 expression. The CIE-induced changes in astrocytic GAT3 expression were paralleled by concordant alterations in GABAergic transmission in the CeA, including increased GABA transmission and the emergence of a GAT3-sensistive tonic GABA current. Despite these changes, manipulating astrocytic expression of GAT3 did not alter somatic withdrawal, dependence-escalated drinking, aversion-resistant drinking, or post-dependent ethanol drinking in male or female rats. Overall, our results suggest that chronic ethanol exposure dysregulates elements of GABAergic transmission in the CeA, including increased GABA release (IPSC frequency) and GABA spillover (tonic current), and that astrocytes play a homeostatic role to oppose excess GABA release by upregulating GAT3 expression and increasing their proximity to the synapse, enabling the GAT3 transporter to provide increased regulation over the tonic GABA current.

Most studies examining the effects of ethanol on astrocytes have focused on changes in gross morphology, including size, shape, and number[78] or broad transcriptomic alterations in astrocytes following ethanol exposure[79, 80]. Although, recent research has begun to explore ethanol effects on astrocyte-neuron interactions[69, 81] and manipulations of astrocyte function on ethanol-related behaviors[82, 83]. The experiments presented in this study describe both effects of ethanol exposure on astrocyte plasticity and functional astrocyte manipulations to understand how ethanol-induced changes in astrocyte structure/function influence behavioral outcomes. Dynamic changes in astrocyte-synapse proximity are observed following exposure to a number of different drugs of abuse[54, 68, 71, 84]. For example, one of the first reports linking changes in structural astrocyte plasticity with behavioral outcomes found that during cued reinstatement to heroin, astrocyte processes rapidly move toward the synaptic space and inhibit heroin seeking behavior[54]. While our results suggest that withdrawal from chronic ethanol exposure induces structural plasticity in astrocytes, promoting insertion of astrocyte processes towards the synapse, future studies are needed to ascertain whether these structural changes have functional implications for the behavioral consequence of ethanol dependence.

Dysregulation of GABA transmission in the CeA is a hallmark of ethanol dependence[6, 70]. Repeated bouts of ethanol exposure increase synaptic GABA release and elevate extrasynaptic GABA levels[6, 10], in addition to a downregulation of inhibitory GABA_B_ auto receptors[13]. Results from the present study are in accordance with these findings as the slice electrophysiology experiments suggest that dependent animals exhibit increased GABA release and tone as measured by increased IPSC frequency and tonic current. In addition, we extend these observations to another component of GABA regulation by showing that chronic ethanol exposure upregulates astrocytic GAT3 expression and that dependent rats are more reliant on GAT3 function to regulate tonic GABA current in the CeA. Prior reports have observed increases in GAT3 protein expression in several brain regions following chronic ethanol administration, suggesting that increases in this GABA transporter might be a common neural adaptation induced by chronic ethanol exposure[85, 86]. The upregulation of GAT3 expression, together with the increase in synaptic proximity, positions astrocytes in an ideal configuration to uptake extrasynaptic GABA and bias GABA diffusion, potentially limiting the impact of GABA spillover on neural circuit function. A similar role for astrocytic GAT3 in gatekeeping GABA levels has been observed in the thalamus and suggests a general role for astrocytes in maintaining neurotransmitter homeostasis[43].

Despite the consistent increase in astrocytic GAT3 levels following chronic ethanol exposure, this adaptation appears largely dispensable for several behavioral phenotypes of dependence. For example, the viral GAT3 manipulation approaches did not alter somatic withdrawal, escalated ethanol drinking, aversion-resistant drinking, or post-dependent drinking in CIE-exposed rats. These results are somewhat consistent with a prior report demonstrating that GAT3 knockdown in the CeA increased GABA tone but did not alter ethanol self-administration[59]. A limitation of the viral gene manipulations is that they constitutively overexpress or knockdown gene expression, which presents the possibility for compensatory adaptations to counteract these manipulations over time. To account for this possibility, we performed acute pharmacological blockade of GAT3 in the CeA of dependent rats. However, this manipulation also had no effect on dependent drinking. Overall, these findings support the theory that astrocytic GAT3 does not regulate ethanol drinking phenotypes associated with ethanol dependence.

An alternative hypothesis for the current findings is that the astrocytic adaptions induced by CIE have a more complex and potentially cell-type specific role within the CeA. The CeA contains well characterized subpopulations of recurrent inhibitory GABAergic microcircuits that regulate diametrically opposed behaviors[87, 88]. For example, CeA neurons that express Corticotropin-releasing factor (CRF+) regulate ethanol drinking in dependent rats[9] and form reciprocal inhibitory connections with somatostatin-expressing CeA neurons[89]. In addition, there is also notable astrocyte heterogeneity within the CeA. Subpopulations of astrocytes that express the oxytocin receptor versus those that do not differentially regulate affective states via their influence on neuronal activity in distinct subregions of the CeA[90]. Moreover, different neuron types in the CeA undergo selective adaptations following chronic ethanol exposure[91, 92]. For example, CeA neurons expressing Corticotropin-releasing factor receptor 1 (CRF1R+) exhibit a loss of tonic GABA current following CIE exposure, whereas CRF1R-neurons begin to display a tonic GABA current following CIE exposure. Moreover, protein kinase C delta (PKCd) expressing neurons in the CeA appear uniquely sensitive to transcriptomic alterations induced by chronic ethanol exposure[79]. Therefore, it is plausible that manipulating astrocytic GAT3 levels throughout the CeA affected multiple cell types differently and potentially neutralized any effect at the behavioral level.

Though we still lack a thorough understanding of how ethanol-induced dysregulation of astrocytes influences neuronal function and behavior, recent progress has provided promising insights[71, 78, 80, 82, 83]. The findings described in the present study demonstrate that CIE exposure remodels astrocyte-neuron interactions in the CeA and suggests that CeA astrocytes may be involved in GABAergic homeostasis during ethanol withdrawal. Molecular substrates underlying dependence have long been targets of medication development, but have yielded negligible results thus far[7, 93–95]. Therefore, exploring non-neuronal cell types might offer an alternative avenue for identifying novel substrates involved in negative adaptations during dependence and facilitate development of improved medications for alcohol use disorder.

## Author contributions

TBN and LJC designed the experiments and co-wrote the manuscript. TBN performed experimental data collection and analysis. ETW, DTV assisted with behavioral experiments. JDO performed slice electrophysiology experiments and analysis, AK and MDS provided training for astrocyte imaging experiments and valuable discussion.

